# Movement patterns of free-roaming dogs on heterogeneous urban landscapes: implications for rabies control

**DOI:** 10.1101/684381

**Authors:** Brinkley Raynor, Micaela De la Puente-León, Andrew Johnson, Elvis Díaz-Espinoza, Michael Z. Levy, Sergio E. Recuenco, Ricardo Castillo-Neyra

## Abstract

In 2015, a case of canine rabies in Arequipa, Peru indicated the re-emergence of rabies virus in the city. Despite mass dog vaccination campaigns across the city and reactive ring vaccination and other control activities around positive cases (e.g. elimination of unowned dogs), the outbreak has spread. Here we explore how the urban landscape of Arequipa affects the movement patterns of free-roaming dogs, the main reservoirs of the rabies virus in the area. We tracked 23 free-roaming dogs using Global Positioning System (GPS) collars. We analyzed the spatio-temporal GPS data using the time- local convex hull method. Dog movement patterns varied across local environments. We found that water channels, an urban feature of Arequipa that are dry most of the year, promote movement. Dogs that used the water channels move further, faster and more directionally than dogs that do not. Our findings suggest that water channels can be used by dogs as ‘highways’ to transverse the city and have the potential to spread disease far beyond the radius of control practices. Control efforts should focus on a robust vaccination campaign attuned to the geography of the city, and not limited to small-scale rings surrounding cases.

## Background

Since March 2015, hundreds of rabid dogs have been detected in the city of Arequipa in southern Peru, following 15 years of epidemiological silence (1,2). The system of water channels in Arequipa, a unique urban landscape feature, has been associated with the location of rabid dogs, suggesting that the city’s complex environment influences the transmission of the disease (3). In addition to the landscape, Arequipa, like much of Latin America, is characterized by complex socio-ecological characteristics that complicate efforts to eliminate rabies.

Dog ownership practices interplay with unique geographical features of Arequipa city to impact both dog ecology and rabies control. In many areas of Latin America, including Arequipa, owned dogs commonly have access to the street without owner supervision (4); these dogs receive varying degrees of feed and veterinary care (5-10). Populations of free-roaming dogs without owners (strays) are believed to be small due in part to the high mortality of stray dogs (11,12).

There is some evidence that Arequipa’s landscape may affect free-roaming dog movement and ecology. A distinctive geographic feature of the city is an extensive system of open water channels that are dry most of the year (Figure 1), and only carry water during the short rainy season between January and March. When dry, these water channels collect garbage and dogs have been observed foraging for food through the waste. These channels have been spatially associated with canine rabies cases (3), and we hypothesize that they might be used as ecological corridors by dogs.

**Figure 1:**
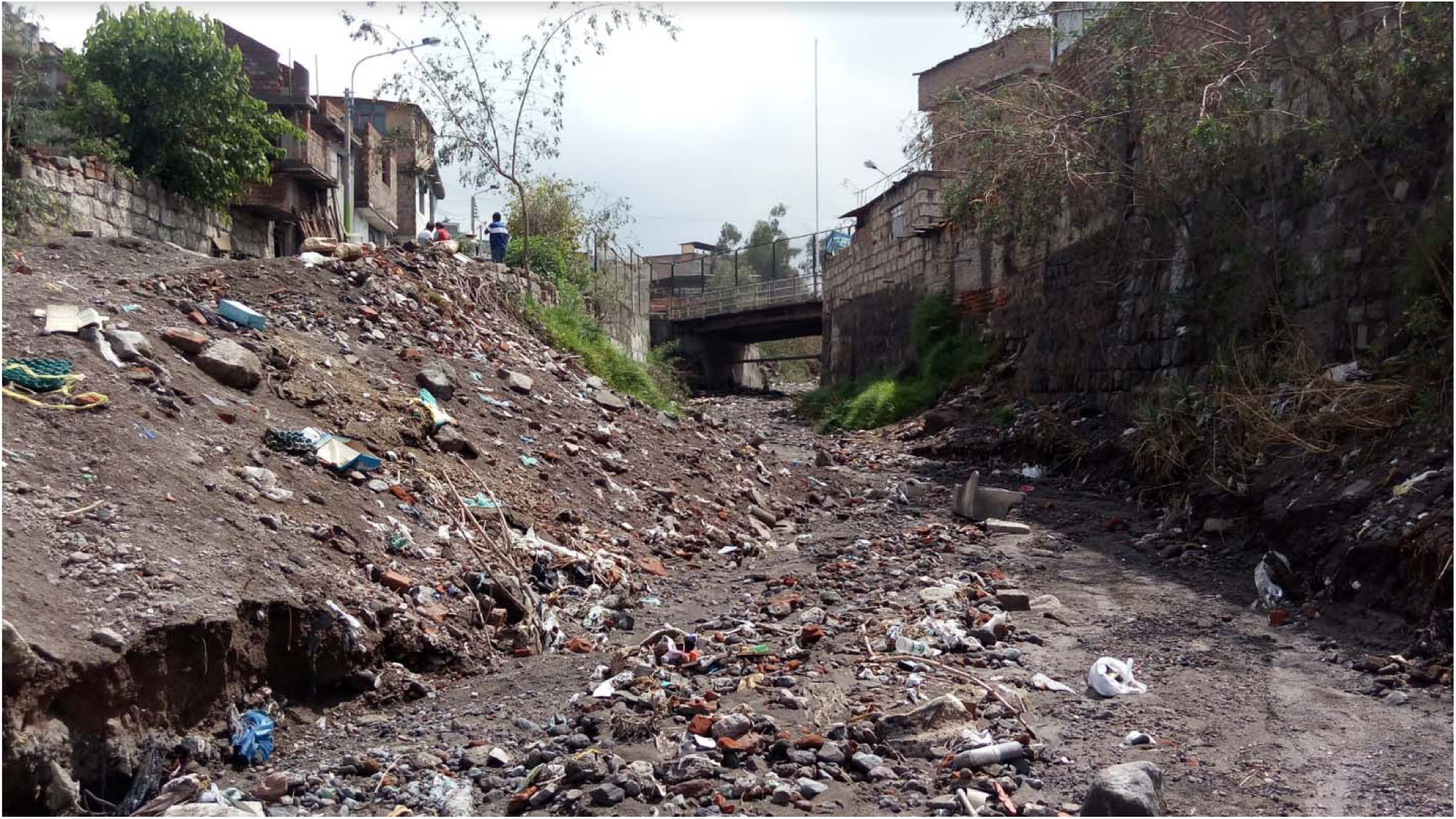
Dry water channels. Water channels traversing the City of Arequipa, Peru are dry most of the year. They accumulate trash and attract free roaming dogs.

The ecology of urban rabies is intimately connected with dog population dynamics and home range, critical concepts in disease ecology (13-15). In a seminal paper introducing home range in 1943, Burt describes the concept of home range as “the area traversed by the individual in its normal activities of food gathering, mating, and caring for young” (15). The study of animal home range has advanced considerably with the incorporation of Global Positioning System (GPS) technology and has allowed for sophisticated analyses of temporal and spatial usage data (16).

There is very little literature on the home ranges and movement patterns of free-roaming dogs; most published studies are focused on rural settings (17-21). These studies in rural areas suggest that dogs home ranges are heterogeneous with variation being influenced by dog function, resource availability, biological characteristics and human interactions (17-21). In Chile, rural dogs had a preference to move using trails and roads, avoiding dense vegetation (21). Very little is known about how the urban landscape affects the space use and movement of free-roaming dogs in cities. A study in Baltimore, Maryland done before the application of GPS technology found that dogs used alleys behind houses to forage and move (22). The larger set of studies on wild animals in urban environments also suggests that animal movement is strongly influenced by landscape features. For instance, cougars preferentially use dry river beds to move in bordering urban areas in California, foxes use ravines in Toronto to move across the city, and bobcats and coyotes use culverts and connected fragments of vegetation to move through southern California (23-25).

Since the outbreak began in March 2015 to August 2019, local authorities have identified more than 160 rabid dogs in 11 out of the 14 districts of Arequipa (26-29). The current response protocol is to conduct ring containment activities, which include vaccination of owned dogs and culling of unowned, and sometimes owned, free-roaming dogs around each detected case (30). However, these focalized small-scale interventions are not supported by data or scientific evidence (31,32). Small, reactive vaccination campaigns are also not recommended when the disease has spread over extended areas (31) and euthanasia is recommended only for rabid dogs or suspected cases. A deep understanding of dog ecology is critical to rabies control (14) and elucidating how dogs move within cities will provide important insights towards this end. In this study we focused on the movements of free-roaming owned dogs. Our objectives were to quantify the variability in dogs’ home ranges, to characterize dogs’ movement in urban areas, and to identify how water channels might affect them.

## Methods

### Ethics

This study was reviewed and approved by the Institutional Animal Care and Use Committee of the Universidad Peruana Cayetano Heredia (approval number 67258).

### Study Sites and Population

The city of Arequipa is surrounded by a desert. There are some wild mammals in this desert, but they are few and far between, and no evidence that wildlife species sustain, or could sustain, rabies virus transmission (3). Three urban localities, a political city district subdivision, were selected for inclusion in this study (Figure 2). One of the localities is urban and intersected by a water channel; the other two localities are peri-urban, one intersected by a water channel and one approximately a kilometer from the nearest water channel. Houses were selected purposively based on location with respect to the water channels. Dogs were eligible for inclusion if they were at least 1 year old, were apparently healthy at the time of the first visit, and were medium or large dogs (at least 35cm at the withers)(33). When multiple dogs were found in a house, we chose the first dog seen by the field team. Dogs of both sexes were included. The characteristics of the selected dog population are summarized in Table 1.

**Figure 2:**
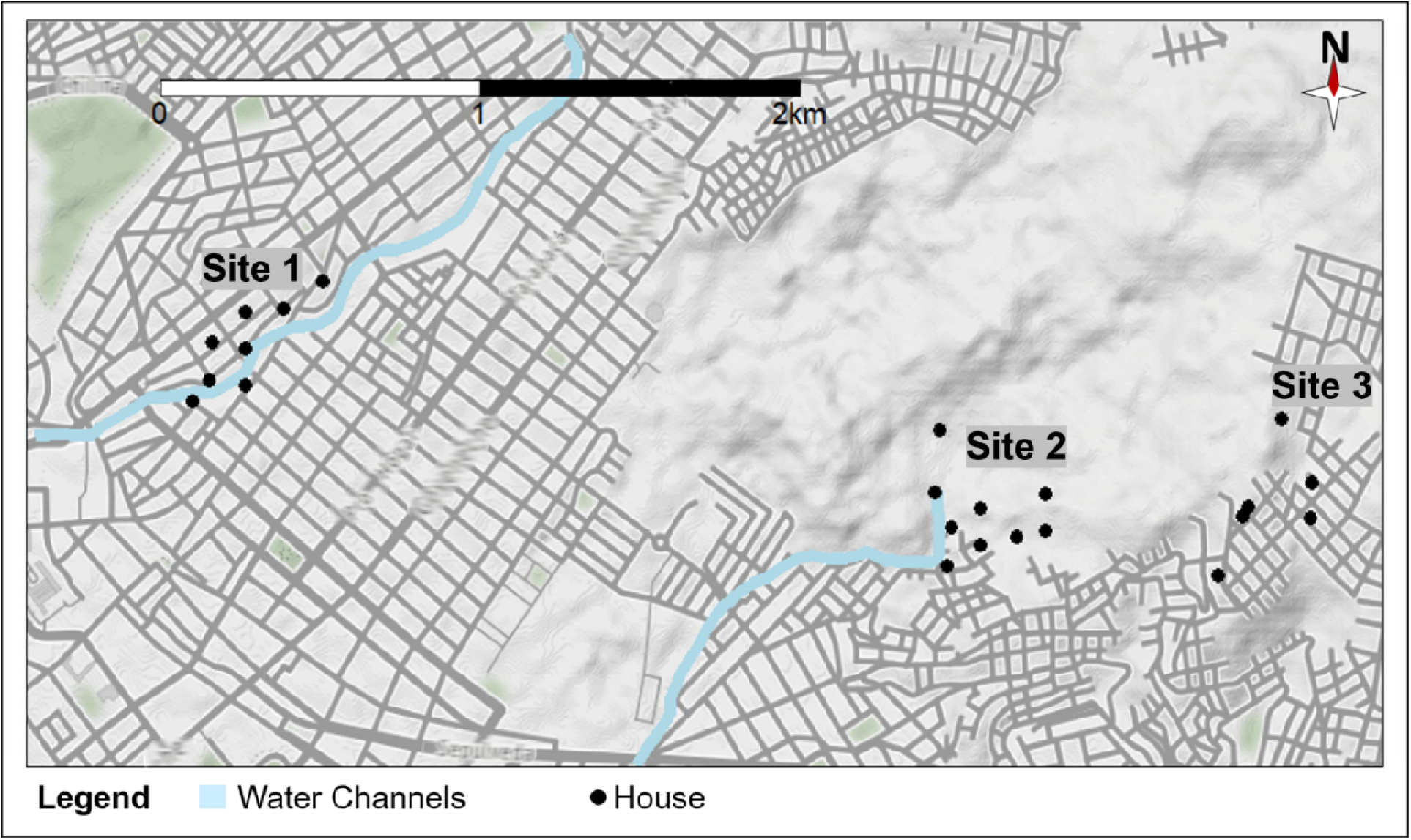
Map of study area and location of dogs’ houses. Dogs were selected for inclusion in the study in both urban and peri-urban locals with houses either directly adjacen to a water channel or approximately a kilometer from a water channel.

**Table 1:** Dog characteristics of study population.

|  |  | **Urban near water channels (n=8)** | **Periurban near water channels (n=9)** | **Periurban away from water channels (n=6)** |
| --- | --- | --- | --- | --- |
| **Sex [no. (%)]** | Male | 6 (75.0%) | 7 (77.8%) | 6 (100.0%) |
|  | Female | 2 (25.0%) | 2 (22.2%) | 0 (0.0%) |
| **Age [no. (%)]** | <= 3 years | 5 (62.5%) | 4 (44.4%) | 0 (0.0%) |
|  | 4-6 years | 2 (25.0%) | 4 (44.4%) | 4 (66.6%) |
|  | 7-9 years | 0 (0.0%) | 0 (0.0%) | 2 (33.3%) |
|  | > 9 years | 1 (12.5%) | 1 (11.1%) | 0 (0.0%) |

### GPS tracking of dogs

To record dog locations and speeds we used IgotU® GT-120 (Mobile Action Technology) GPS loggers that have shown good accuracy in urban environments in Peru (point accuracy of 4.4 m and line accuracy of 10.3 m) (34) and have been used to track free-roaming domestic animals (35). Given the high-speed dogs can achieve, we programmed the GPS receivers to log the dog’s geographical coordinates every 3 minutes. For every dog, o was turned on, and placed in a nylon collar on the dog (Figure 3). After 4 to 7 days, depending on dog and owner availability, we returned to enrolled households. If the dog owner allowed us to continue tracking the dog, we changed the GPS receiver in the collar for another one with fresh batteries or took the collar off if the dog owner requested to drop from the study. Follow-up times per dog varied from 4 days to 4 weeks. We estimated the correlation between observation period duration and home range to evaluate if home range estimates were affected by the variability in observation duration.

**Figure 3:**
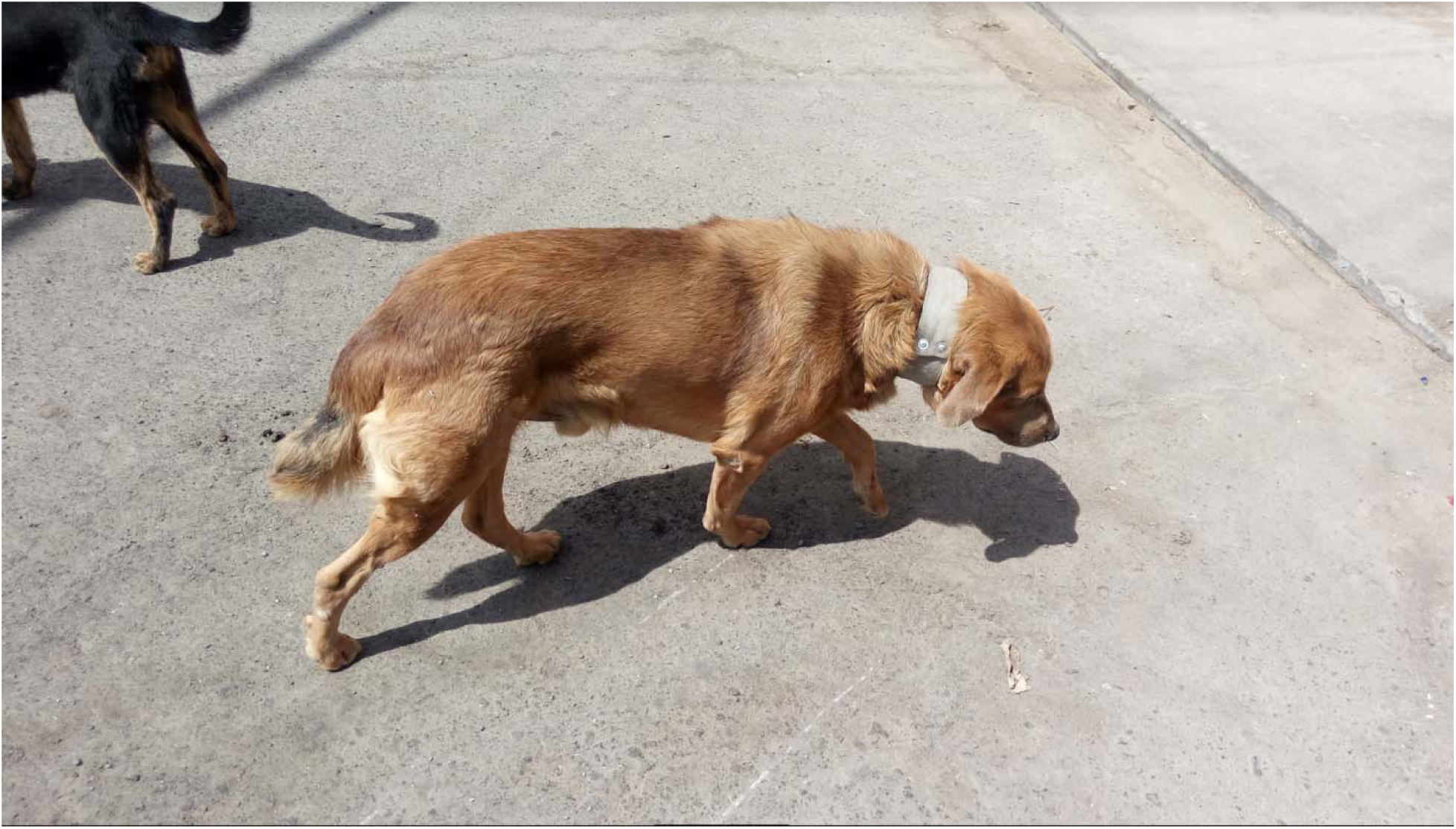
Dog wearing GPS collar. Dog fit with GPS collar, IgotU® GT-120 (Mobile Action Technology) GPS loggers. Battery life of the collars lasted 4-7 days while recording GPS coordinates every 3 minutes.

### Data cleaning

In cities there is potential for urban structures to affect the accuracy of GPS loggers. To handle this potential issue, we excluded speed points that were higher than 30% the fastest speed recorded for a dog (17.003 m/s) (36). This speed limit (5.101 m/s) is similar to the speed limit used by Dürr and Ward (5.556 m/s) in their study of Australian free-roaming rural dogs (18).

## Statistical Analysis and Mapping

### Time spent in the water channels

We used the proportion of recorded locations within 30 m of the water channels as a proxy for the proportion of time spent within these structures. We compared the time spent in the water channels between dogs living close and far away from the water channels. We eliminated from this analysis the points recorded at the dogs’ house, so that the proportion of time not spent at the water channel represents time spent on the streets or other open areas (e.g. parks).

### T-LoCoH

We analyzed dog home ranges and other dog movement metrics using the Time Localized Convex Hulls (T-LoCoH) method. T-LoCoH extends the classically used Local Convex Hull (LoCoH) to include temporal data in the calculations. Briefly, the LoCoH method creates individual convex polygon hulls from a set of selected nearest neighbors around each GPS point without taking account when those nearest neighbors were registered (37). T-LoCoH goes beyond the classic LoCoH by using “time-scaled distance (TSD)” instead of geographical distance to create local hulls around GPS points (16). This approach is ideal for modern GPS data that standardly includes a time stamp with the GPS coordinates. The advantage of T-LoCoH to LoCoH and other previous methods is that nearest neighbors are selected based on closeness in both space and time with the weight given to time closeness based on parameters that can be inferred from the data (16).

### Parameter estimation

We used the *a*-method of nearest neighbor selection. The *a*-method selects neighbors by finding the difference in time scaled distance between the parent point and other points in the set (16). Then the nearest points are added in ascending order of time scaled distance until some value, *a*, is reached (16).

In order to calculate time scaled distance, the central feature of T-LoCoH, a scaling factor, *s*, is needed. The *s* value determines the weight that time is given when selecting nearest neighbors, where *s*=0 would not take time into account when selecting neighbors, while a *s*=1 would select neighbors completely based off of time (16). We chose our *s* value based off temporal cyclicity observed in the field. The free-roaming dogs of Arequipa exhibited distinct morning, afternoon, evening, and night behaviors, so we decided to base our *s* value off a 6-hour cycle time of interest.

For each dog we estimated an *s* value such that the number of time-selected and space-selected neighbors were balanced equally for specified observation periods of interest (38). Once we calculated an *s* value for each dog, we calculated the *a* values. To do this, we selected an *a* value that did not cause drastic increases or decreases in isopleth area and isopleth perimeter (38). A local convex polygon was drawn around each set to create a hull and a combined set of hulls sorted in a specific way is referred to as an isopleth (16).

### Spatio-temporal Models

We used the T-LoCoH package in R (16) to create hulls; the distance between points was calculated in time-scaled distance using our selected *s* value and then points included in each hull were found by summing the distance between closest neighbors until the *a* value was reached (but not surpassed) (38). Isopleths were sorted by both density of points and by eccentricity of the minimal area bounding ellipse eccentricity of each hull (16). Eccentricity is a measure of how directed a movement is; very directed movements having an eccentricity close to 1, while the eccentricity of more random movements is closer to 0 (16). Core home ranges were calculated using the densest 50% isopleth, while extended home range the densest 95% isopleth (18,39).

### Mapping

Dog home ranges were mapped using the R package ggmap (40) on top of a background generated from OpenStreetMap data (41).

### Statistics

All analyses, maps, and graphs were produced with R (42). We compared the core (50% isopleth) and extended (95% isopleth) home ranges, farthest GPS point recorded from the dogs’ homes, and the average eccentricity of hulls between dogs living in 3 different areas and between dogs exhibiting different behaviors in respect to water channel usage. To group dogs based on water channel usage, first a group was made of dogs who had no GPS locations recorded in the water channels. Then the total length utilized (length of water channel in between farthest points in the water channel recorded) was calculated for all remaining dogs. K-means analysis was used to separate dogs into light and heavy water channel users.

## Results

We were able to collar 25 dogs and retrieve data from 23 of them. One dog did not have the collar at the second visit, and we decided to remove it from the study. For another, the logger did not record any data, and the dog owner did not want to continue in the study. The sex and age of the 23 dogs are summarized in Table 1. One dog was included in the study because the owner said it was one year old, but later she rectified it was 10 months old. We kept the dog in the study. Spaying and neutering dogs is a very rare practice in the study area and, as expected, none of the dogs tracked in this study was spayed or neutered.

After data cleaning, our total data set of 23 dogs included 74,120 observations. 595 observations were excluded for being faster than our set cutoff speed. We suspect that most points recorded above our cutoff speed were due to GPS error. The excluded points contained some few speeds that could plausibly be attributed to true dog movement.

We observed high variability in home ranges (Figure 4) as well as eccentricity (Figure 5). We also found high variability among what times of the day dogs moved the most (Figure 6). We found important differences when comparing dogs based on their water channel usage. In Figure 5 and 6 we show the home ranges and eccentricity maps of 3 dogs, A, B, and C. These dogs exemplify movement patterns of dogs that either never use the water channels (A), lightly use the water channels (B) or heavily use the water channels (C).

**Figure 4:**
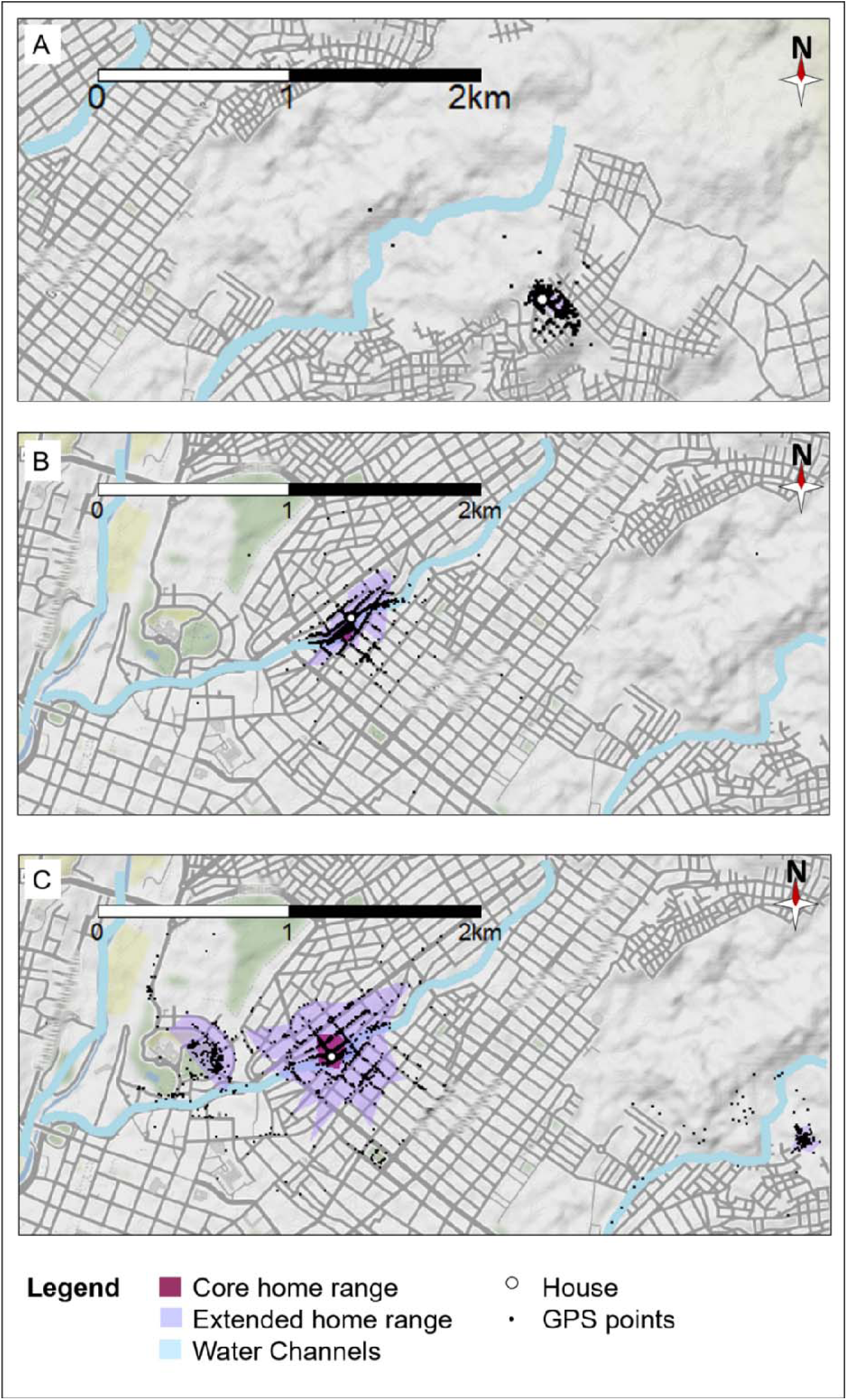
Core and extended home range of 3 dogs that exemplify different movement behavior patterns. Maps of the core (50% isopleth) and extended (95% isopleths) home ranges for dogs with three different behavior patterns. Dog A tends to stay at home or close to the house most of the day and never goes into the water channels. Dog B stays in a small, defined area around her house and goes into the water channel in proximity to her house. Dog C ranges far from his house and has points going along multiple water channels.

**Figure 5:**
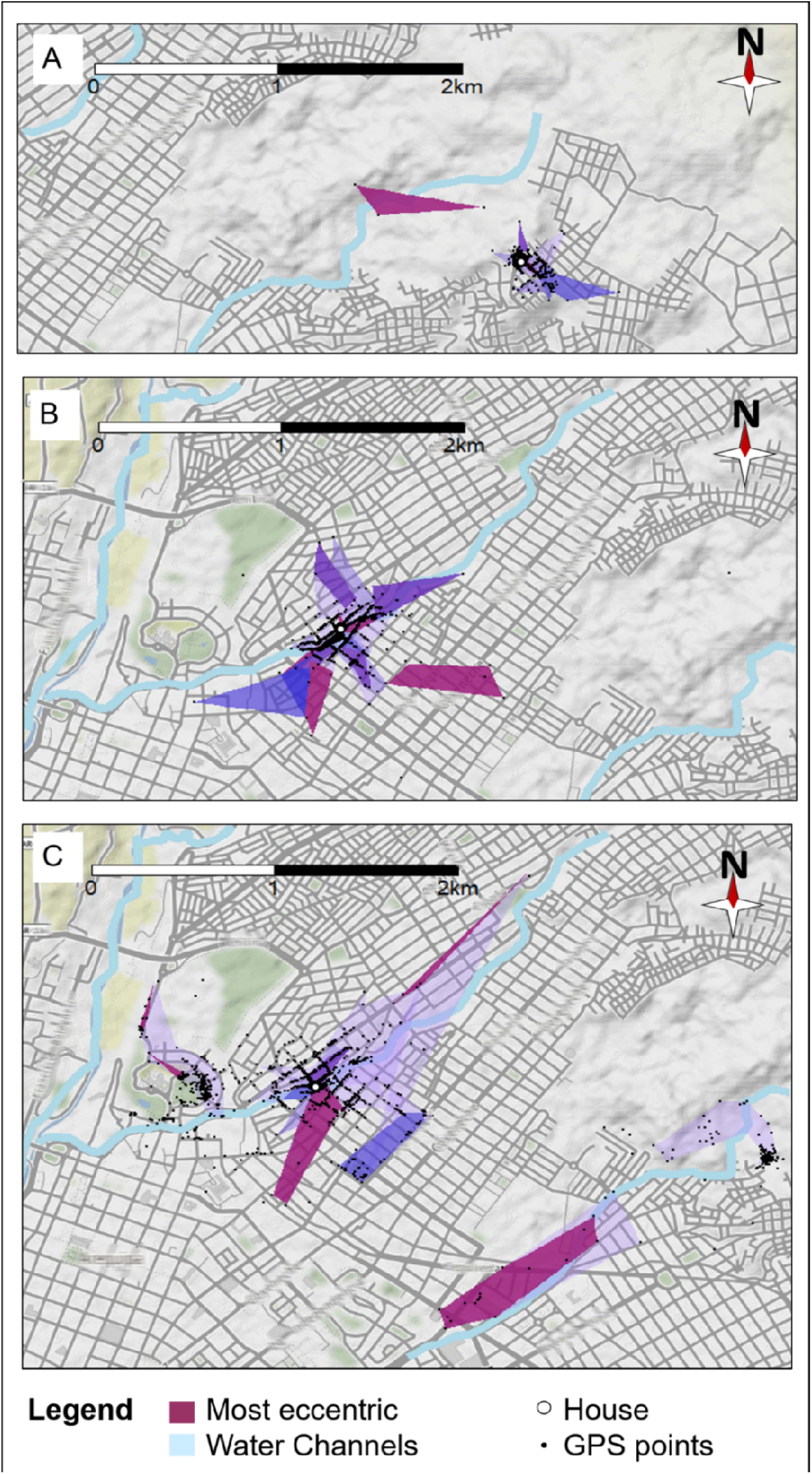
Eccentricity of 3 different dogs exemplifying different movement behavior patterns. Maps of Dog A, B, and C’s isopleths sorted by eccentricity: the redder the isopleth color, the greater eccentricity values of the isopleths.

**Figure 6:**
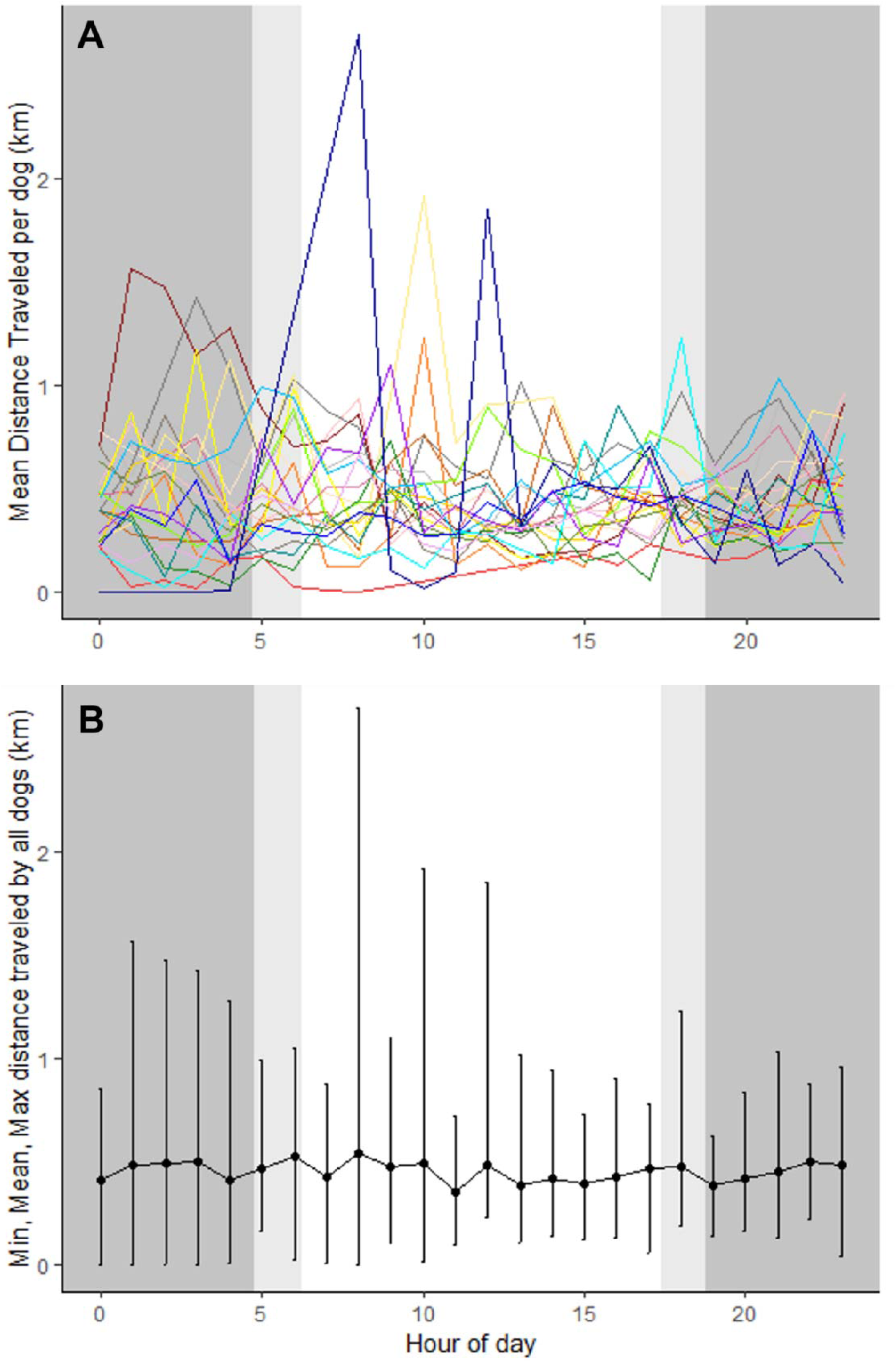
Average distance travelled per hour of day by dogs in the city of Arequipa, 2018. The background colors represent nighttime (dark grey), dawn/dusk (light grey), and daytime hours (white). Panel A is a plot of the mean of each individual dogs’ distance traveled for each hour of the day. Panel B represents the mean of the results shown in panel A with the range bars displaying the minimum and maximum.

Dogs that used the water channels more tended to have the largest core and extended home ranges. Core home ranges, based on the 50% density isopleth, ranged from 0.00013 km^2^ to 0.46 km^2^. Extended home ranges, based on the 95% density isopleth, ranged from 0.0012 km^2^ to 3.70 km^2^. Water channel usage was also associated with moving the longest distance from home (p=0.002) and moving with higher directionality (p=0.027) (Table 2). One dog that regularly used the water channels traveled up to 14 km from its home. The maximum speed achieved by dogs also increased with water channel usage, but there was not statistical association. Interestingly, no significant differences in dog movement patterns were found in dogs grouped by home location.

**Table 2:** Area utilization based on dog’s usage of water channel.

|  | Heavy water<br>channel usage<br>(n=4) | Light water<br>channel usage<br>(n=13) | No water channel<br>usage<br>(n=6) | p-value (Kruskal<br>Wallis) |
| --- | --- | --- | --- | --- |
| **Core home range**<br>**area (50%**<br>**isopleth) in km<sup>2</sup>** |  |  |  |  |
| Median (IQR) | 0.156 (.280) | 0.011 (0.015) | 0.003 (0.006) | 0.010 |
| mean (SD) | 0.199 (0.208) | 0.013 (0.011) | 0.011 (0.019) |  |
| **Extended home**<br>**range area (95%**<br>**isopleth) in km<sup>2</sup>** |  |  |  |  |
| Median (IQR) | 1.83 (1.83) | 0.055 (0.065) | 0.017 (0.018) | 0.002 |
| Mean (SD) | 1.951 (1.447) | 0.102 (0.113) | 0.020 (0.018) |  |
| **Longest distance from home** |  |  |  |  |
| Median (IQR) | 3.572 (3.533) | 1.355 (1.629) | 0.982 (0.291) | 0.002 |
| Mean (SD) | 6.043 (SD) | 2.251 (SD) | 0.866 (SD) |  |
| **Eccentricity** |  |  |  |  |
| Median (IQR) | 0.771 (0.037) | 0.618 (0.079) | 0.613 (0.057) | 0.027 |
| Mean (SD) | 0.746 (0.055) | 0.624 (0.078) | 0.622 (0.037) |  |
| **Maximum speeds** |  |  |  |  |
| Median (IQR) | 4.994 (0.057) | 4.772 (0.511) | 3.856 (0.882) | 0.197 |
| Mean (SD) | 4.971 (0.091) | 4.642 (0.495) | 3.734 (1.462) |  |

## Discussion

We found a strong effect of the urban landscape on dog movement in Arequipa, Peru. Dogs that spend more time in the water channels have more linear movements, significantly larger home ranges, and venture further from home than those that spend little or no time in these channels. Our findings suggest that the water channels in Arequipa function as ecological corridors. These corridors might greatly complicate the control of the transmission of rabies.

The traditional response to every detected rabid dog is to conduct “ring containment”. The principle of ring containment is adapted from ring vaccination where all contacts with a positive case are immunized to create a buffer preventing disease spread (43). In Peru, ring containment consists of visiting an area of a determined radius (3 to 5 city blocks in Arequipa) around the location where the rabies positive dog was found to vaccinate unexposed owned dogs, to eliminate stray dogs and exposed or potentially exposed owned dogs, and to simultaneously conduct health promotion focused on rabies and find people who might need post-exposure prophylaxis (30). The ring containment area for each case seems to be dictated by logistics, mostly how many personnel are available in the zone. We found that apparently healthy dogs move far beyond the current fixed-radius (300 to 500 m depending on personnel) ring containment on a regular basis. Rabid dogs usually exhibit erratic behavior with some records of dogs moving more than 15 km from home (44), therefore, it is unlikely that small-scale ring containment activities would reach all or most dogs that may have come into contact with a rabid individual. It is not feasible to increase the radius of the preventative ring activities when the rabies control teams are frequently understaffed. In a door-to-door survey conducted in the same study area, 25% of owned dogs have unrestricted access to the streets (4). Under these conditions, implementing dog-centered small-scale activities that might include culling and injection-delivered vaccination becomes challenging due to difficulty of finding dogs at home and distinguishing between free-roaming owned dogs and strays.

Water channels and other similar structures that function as ecological corridors have implications for appropriate modelling of disease spread (25,45). Infected animals moving directionally along these corridors have the potential to connect parts of an urban landscape that are not geographically contiguous or even proximate. The increased connectivity created by the city landscape has implications on vaccination goals. It has been reported that small pockets of unvaccinated dogs can sustain rabies transmission, (46), and these ecological corridor have the potential to connect otherwise separated sub-optimally vaccinated populations increasing the chances of extending the area of the epidemic (47,48).

By tracking apparently healthy dogs, we have gained insight on the impact of the urban landscape of movement of owned free-roaming dogs, an important reservoir or rabies virus. It is known that rabies can change the movement patterns of dogs (49), However, specific dogs that have movement patterns that put them in contact with many other dogs have a greater likeliness of virus transmission (50), therefore, it is as important to understand how uninfected dogs move. Our study captured the movements of 23 dogs; a larger follow up study is needed to obtain reliable parameters to model rabies transmission in urban landscapes that favor long, directed and fast incursion of dogs or packs of dogs into new areas. Finally, our categorization of water channel usage includes inherent bias with increased water channel usage being related to increased home range. It is possible that a larger study with more followed up individuals would allow the use of synthetic likelihoods (51) to tease apart the effect of water channels on these long, directed “flights”.

Interestingly, we found significant movement at night for over half the dogs tracked. These observations, contradict the paradigm that owned free roaming dogs have a more diurnal pattern while stray dogs exhibit a more nocturnal pattern (22), and advise caution when following guidelines that state that daytime counting methods to estimate the dog population are appropriate for owned dogs (52). Even though we did not track stray dogs to compare against them, our data does not show any trends of owned dogs moving more during daytime, instead varying significantly by individual. Any strategies to control stray dog populations should not focus on dogs that are out at night (as have been suggested by local authorities) as these may be owned free-roaming dogs, not strays. In addition, assessment of the dog population size might be inaccurate if one of the assumptions of the methods is that most owned dogs are diurnal.

Creating a successful dog rabies control program and transforming it into a sustainable prevention program requires planning with knowledge of epidemiological concepts and deep understanding of local populations and local needs (14,53). Around the world, dog rabies control programs have demonstrated that regional canine rabies elimination is feasible (5,54-56). Particularly, in Latin America the control programs have been very successful reducing the burden of disease significantly (5,56). Our findings with the water channels reinforce the importance of focusing on city-wide approaches with vaccination programs that reach both, optimal levels and even coverage across the city. Our findings, and previous findings of rabid dogs spatially associated with these water channels, suggest that surveillance activities in and around these structures, as well as vaccinating the animals that move along them (most feasibly with an oral vaccine) could be critical to re-eliminating rabies virus from the city.

## Acknowledgements

Dr. Seth O’Neill and the Center for Global Health, Tumbes – UPCH for lending some of the GPS loggers.

Lina Mollenasa, Liz Leon, Edwin Cardenas, and Julio Meza for their hard work in the field handling and collaring the dogs.

© OpenStreetMap contributors for data used to generate map figures

